# Alternative splicing across the *C. elegans* nervous system

**DOI:** 10.1101/2024.05.16.594567

**Authors:** Alexis Weinreb, Erdem Varol, Alec Barrett, Rebecca M. McWhirter, Seth R. Taylor, Isabel Courtney, Manasa Basavaraju, Abigail Poff, John A. Tipps, Becca Collings, The CeNGEN Consortium, Smita Krishnaswamy, David M. Miller, Marc Hammarlund

## Abstract

Alternative splicing is a key mechanism that shapes neuronal transcriptomes, helping to define neuronal identity and modulate function. Here, we present an atlas of alternative splicing across the nervous system of *Caenorhabditis elegans*. Our analysis identifies novel alternative splicing in key neuronal genes such as *unc-40*/DCC and *sax-3*/ROBO. Globally, we delineate patterns of differential alternative splicing in almost 2,000 genes, and estimate that a quarter of neuronal genes undergo differential splicing. We introduce a web interface for examination of splicing patterns across neuron types. We explore the relationship between neuron type and splicing patterns, and between splicing patterns and differential gene expression. We identify RNA features that correlate with differential alternative splicing, and describe the enrichment of microexons. Finally, we compute a splicing regulatory network that can be used to generate hypotheses on the regulation and targets of alternative splicing in neurons.

## Introduction

Differential alternative splicing is a fundamental mechanism that elevates molecular diversity. Splicing involves processing of pre-mRNAs by the spliceosome, resulting in removal of intronic sequences. Most metazoan genes undergo splicing, and splicing is critical not only for producing mature mRNAs but also for nuclear export and therefore translation. Alternative splicing (AS) occurs when a pre-mRNA is processed in more than one way, resulting in removal of different introns and the consequent production of mature mRNAs with different sequences. AS can alter the mRNA coding potential, resulting in expression of different protein isoforms. AS can also affect the stability and other features of the mRNA itself. The vast majority of human genes undergo AS, and defects in AS have been linked to various diseases ^1^.

Differential AS occurs when AS is regulated spatially or temporally, so that different cells express separate isoforms (differential AS is often referred to simply as ‘AS’; here, the terms are distinct.) In the nervous system, where it is most prevalent ^2, 3^, differential AS controls multiple aspects of neuron identity ^4^, including global AS switches during development ^5^, isoform differences in neuron type specification ^6, 7^, axon specification, guidance, and synaptogenesis ^8, 9^, and has been linked to several brain disorders ^1, 10, 11^. Differential AS has also been studied in cancer; indeed, some forms of cancer are dependent on differential AS ^12^. AS, because it is not cell-specific, can be identified by methods such as sequencing bulk cDNA. By contrast, identification of differential AS requires comparison of splicing between different samples, and thus has been less well characterized. One theme that has emerged is that differential AS is often not all-or-none, but rather is characterized by different ratios of splice isoforms across time or space.

How is splicing regulated? For AS, cis-acting factors—sequences or structures within the pre-mRNA—can regulate splicing diversity. The logic of cis regulation has been analyzed to identify features that give rise to AS ^13, 14, 15, 16, 17^. But for *differential* AS, features within the nascent transcript are not sufficient, since the transcript is the same in all cell types. There must also exist trans-acting factors that regulate AS in a cell type-specific manner, for example by interacting with the spliceosome to promote or inhibit particular splicing events ^1^. Several approaches have been developed to establish splicing regulatory networks ^3, 18, 19, 20, 21, 22, 23^, or to integrate *trans* features within a framework developed on sequence motifs ^24, 25, 26, 27^. Nevertheless, our understanding of the ‘splicing code’—the regulatory framework involving cis and trans elements that determines differential AS across all transcripts—is incomplete.

The nematode *Caenorhabditis elegans* is a powerful model organism for studies of the nervous system including AS ^28^. Although gene expression atlases of developing and aging *C. elegans* cells are now available ^29, 30, 31, 32, 33, 34, 35, 36^, less has been done to systematically establish AS patterns ^37, 38, 39^.

Here, we analyze data generated by the CeNGEN project to produce an atlas of AS for 55 single neuron types in *C. elegans*. We develop analytical tools to make the data available to the research community. We study differential AS between neuron types and show that key neuronal genes display broad patterns of differential AS. Focusing on canonical AS events, we establish overall patterns of differential splicing. Finally, we develop a principled computational approach to extract a regulatory network for differential AS, and use the network to identify candidate factors that regulate differential alternative splicing.

## Results

### Visualization and analysis of alternative exon usage

The CeNGEN project generated a data set covering 55 individual neuron types suitable for splicing analysis. As described previously ^40, 41^, a series of *C. elegans* strains were used, each with specific promoters that uniquely label individual neuron types. For each neuron type, neurons were recovered by Fluorescence Activated Cell Sorting (FACS) from L4 hermaphrodites, with multiple independent biological replicates (average 3.8 replicate per neuron type). Libraries were prepared using an optimized ribodepletion protocol ^41^ and sequenced on the Illumina platform, an approach that yielded robust coverage across the gene body (Fig. 1A). The *C. elegans* nervous system contains 118 canonical neuronal types ^42^. Our recent analysis described 128 neuron types defined by their transcriptome ^32^. Our sorting approach here focused on canonical types, with the exception of the subclasses ASEL and ASER, which were sorted separately. Of the potential 119 types using this approach, 55 types, or 46%, are described in this work. To analyze alternative splicing (AS) in this data set we used three parallel approaches: raw data visualization, local quantification, and transcript-level quantification. We illustrate our approaches on the gene *ric-4*, the homolog of SNAP25, which displays an alternative first exon expressed differentially between neuron types ^43, 44^.

**Figure 1:**
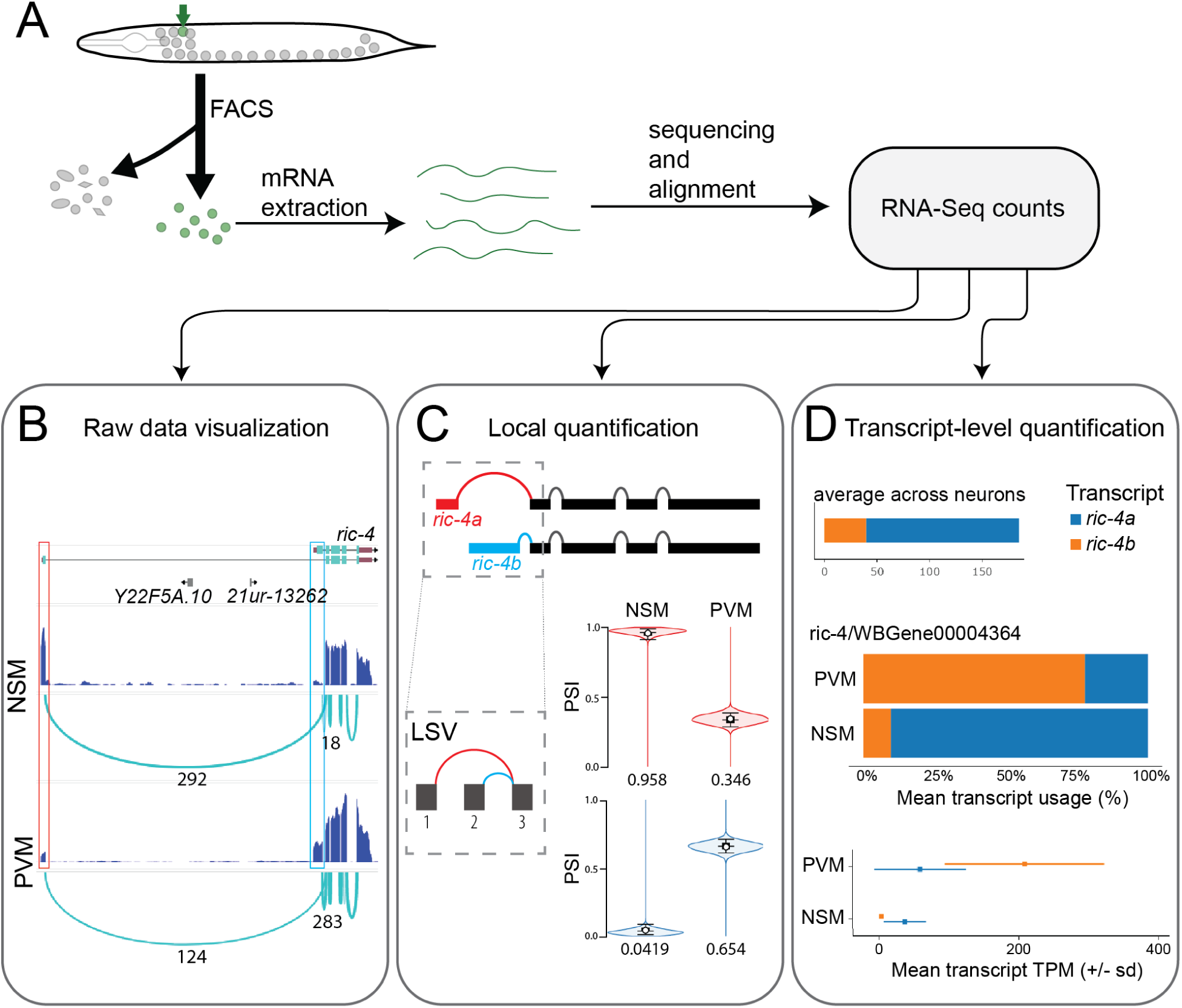
Overview of data collection and splicing analysis. **A:** Schematic of the experimental procedure. **B-D:** Three methods to analyze alternative splicing, applied to the gene ric- 4/SNAP25 in the neurons NSM and PVM. **B:** Raw data visualization. Top track: gene model of ric-4 (along with non-coding RNAs 21ur-13262 and Y22F5A.10). Bottom tracks: Read coverage and junction counts. Numbers denote junction-spanning reads for the splice junctions of interest. Red and blue boxes indicate alternative first exons. **C:** Local Splicing Variation (LSV) visualization. Top: Gene model of ric-4, highlighting the alternative first exons corresponding to ric-4a (red) and ric-4b (blue). Bottom: Posterior PSI estimates from MAJIQ displayed as violin plots. Numbers indicate the posterior expected PSI (summing to one in each neuron, as the red and blue junctions are mutually exclusive). **D:** Transcript-level quantification. Top: Average transcript TPM across all sequenced neurons. Middle: Relative transcript usage (in proportion to the total TPM for the gene, in each neuron), in NSM and PVM. Bottom: Absolute transcript quantification, indicating the mean +/- standard deviation of TPM across samples for each transcript in each neuron.

Raw data visualization is a direct approach for splicing analysis that depends on displaying raw read counts in a genome browser. Direct visualization allows inspection of exon and splice usage in the full context of all the data for that gene. Raw data visualization does not use a statistical model and can be applied to individual biological replicates, to pooled data for each individual neuron type, or to all samples grouped together. Our browser is based on JBrowse2 ^45^. For each individual biological replicate, we generated a pair of browser tracks. The tracks underwent minimal filtering for clarity (see Methods).

First, a density plot indicates the number of reads aligned at a particular genomic position (normalized by the total number of reads in that sample and multiplied by one million, yielding Counts Per Million). Second, a splice junction track indicates the number of junction-spanning reads supporting that junction, without any normalization. We computed similar tracks for each neuron type, using all biological replicates for that type. Here, the density histogram represents the mean coverage across replicates at each genomic position (for each base pair). In addition, the junction-spanning reads (see Methods) are summed for each junction, to give a total junction usage track for that neuron. Finally, to allow rapid examination of a genomic locus across many neurons, we generated an additional set of six “global” tracks: the mean coverage (for each genomic position) across all neuron types, the minimum and maximum coverage at each genomic position across all neuron types, and the sum, minimum, and maximum of junction-spanning reads for each splice junction across neuron types. The mean exonic coverage and sum of splice junction tracks enable convenient visualization of an “average” transcript across neuron types. The minimum and maximum tracks facilitate the identification of rare transcripts: if a single neuron type expresses a given exon, it will be apparent in the maximum coverage track; if a single neuron does not express a given exon, it will not appear in the minimum track (and similarly for splice junctions).

For example, for the gene *ric-4* (Fig. 1B), we apply raw data visualization to the neuron types NSM and PVM. Read coverage shows that the distal first exon (red box), corresponding to transcript *ric-4a*, is preferentially expressed in NSM. PVM displays weaker preferential usage of the proximal first exon (blue box), corresponding to transcript *ric-4b*. In addition, NSM displays 292 junction-spanning reads connecting the distal first exon, and only 18 junction-spanning reads connecting the proximal first exon. By contrast, in PVM there are 124 junction-spanning reads connecting the distal first exon, and 283 junction-spanning reads connecting the proximal first exon. This analysis indicates that *ric-4* undergoes differential alternative splicing.

Although inspecting raw data on the genome browser is useful for visualizing differential AS at the single-gene level, additional methods are needed for genome-wide analysis. For this purpose, we quantified splice junction usage with the software package MAJIQ ^46^. MAJIQ defines Local Splicing Variation (LSV) as splice junctions (SJ) starting from the same source exon or ending in the same target exon. For each LSV, the relative usage of each possible SJ is quantified, and a Percent Selected Index (PSI) is estimated from a Bayesian model. For example, an exon skipping/inclusion event (known as a ‘cassette exon’) is represented by MAJIQ as two LSVs: one upstream LSV, containing two splice junctions (one SJ that links the upstream exon to the cassette exon, and one SJ that skips the cassette to link to the downstream exon), and a second LSV, also with two SJs (one SJ from the upstream exon into the downstream exon, the other SJ from the cassette exon into the downstream exon). Similarly, an alternative first exon is represented in MAJIQ by a single LSV, immediately downstream of the alternative exons. Quantitative data generated using MAGIQ can be represented using VOILA, ^46^, which represents the PSI of junctions belonging to an LSV using violin plots.

In the case of *ric-4*, the alternative first exon is quantified as a single LSV containing two splice junctions, and quantification demonstrates the preferential use of the *ric-4a* exon in NSM and the *ric-4b* exon in PVM (Fig. 1C). We use the quantitative data generated by MAJIQ to analyze global splicing patterns across genes and neuron types in the following sections.

Besides quantifying individual splicing events, it is also useful to visualize DAS in the context of complete transcripts. To address this question, we used the software package StringTie in quantification mode to analyze transcript levels in individual neuron types ^47^. Given a set of annotated transcripts (we used all transcripts annotated in WormBase), StringTie uses a maximum flow computational approach to estimate the expression level of each transcript. Thus, StringTie output represents not only the relative abundance of each transcript in each neuron type, but also their absolute levels (in Transcript Per Million or TPM units).

For example, in the case of *ric-4* (Fig. 1D), StringTie analysis compares the levels of the transcripts *ric-4a* and *ric-4b* in NSM and PVM. First, the transcript *ric-4a* is more common than *ric-4b* when averaging across all neurons. Second, examining the relative transcript usage, 90% of the *ric-4* expression in NSM is attributed to *ric-4a*, whereas 80% of the *ric-4* expression in PVM is attributed to *ric-4b*. Finally, comparing the absolute transcript expression values, *ric-4a* is expressed at 40 TPM in NSM and 60 TPM in PVM, whereas *ric-4b* is expressed at 4 TPM in NSM and 212 TPM in PVM.

Next, we asked if our analysis aligns with previously described alternative exon usage in *C. elegans* neurons. A two-color splicing reporter revealed that *elp-1*/EMAP undergoes differential alternative splicing, with exon 5 skipped in touch neurons ^48^. Similarly, we observe exon 5 skipping in the AVM touch neurons (Fig. 2A). In another case, a cassette exon (11.5) in *daf-2*/IGFR was reported to undergo differential alternative splicing in many neuron classes ^49^ (Fig. 2B). Using local quantification (Fig. 2C), we also find differential alternative splicing at exon 11.5, with the splicing patterns we observe in good agreement with previous results. A similar pattern is seen by visual exploration of the raw data (Fig. 2D, red box). It is interesting to observe that despite relatively clear data for exon 11.5, our transcript-level analysis shows that the transcript known to contain this exon (*daf-2c)* is only modestly enriched in individual neuron types, likely owing to the relatively large number of alternative transcripts of the *daf-2* gene ^43^ (Fig. 2E). Together, these results indicate that our data are consistent with existing *in vivo* observations at single-neuron type resolution.

**Figure 2:**
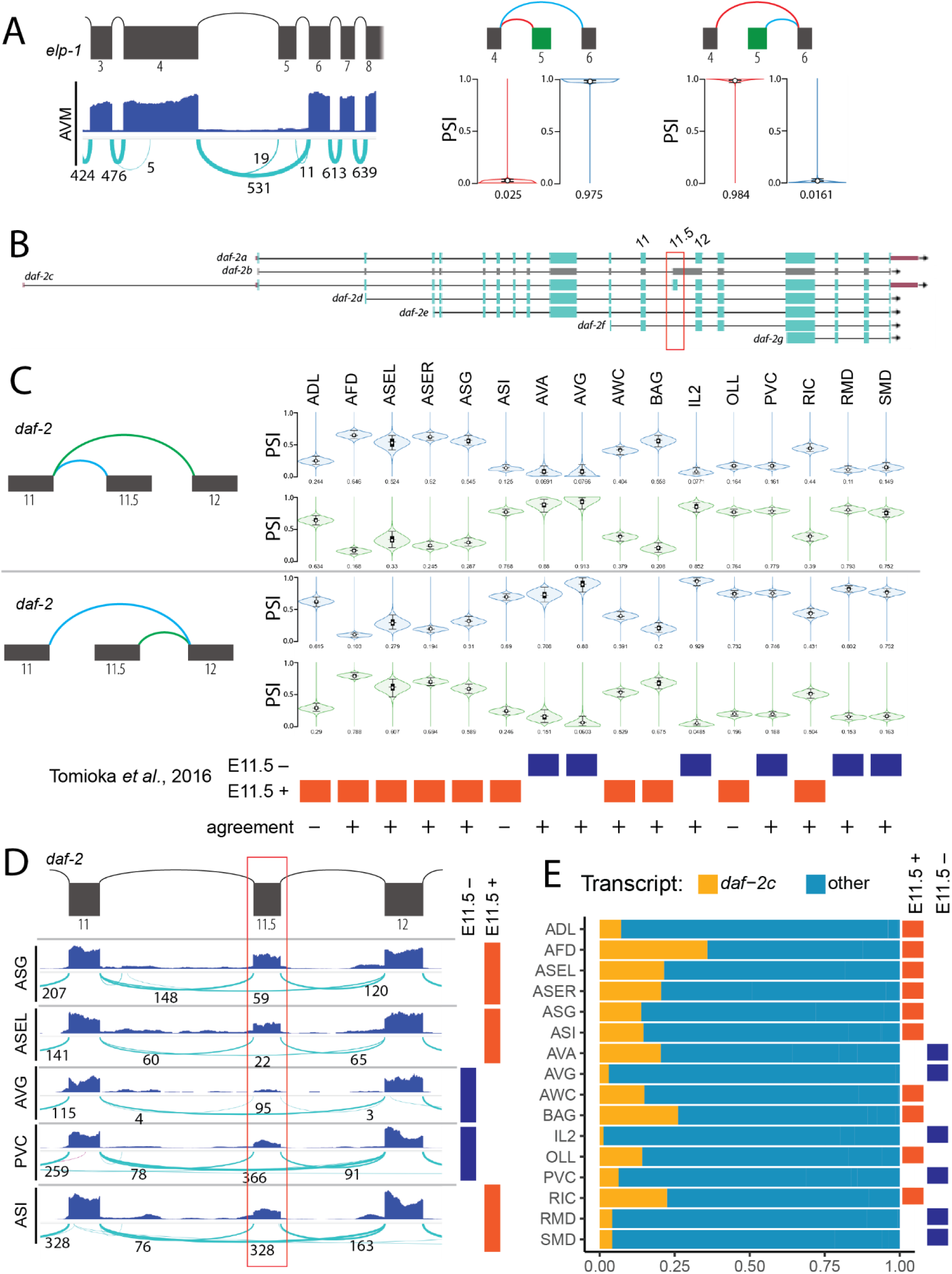
Comparison to previous splicing data. **A:** Exon 5 skipping in elp-1 in AVM neuron. Left: Raw data visualization in the neuron AVM. Right: Local quantification of the two LSVs corresponding to the cassette exon. **B-D:** Cassette exon 11.5 in daf-2 in individual neuron types. **B:** daf-2 gene structure. The cassette exon (red box) is unique to the gene model for daf-2c. **C:** Local quantification of the upstream (top) and downstream (middle) events flanking the cassette exon 11.5. Left: Schematic representation of splice junctions constituting the local event. Right: Splice junction quantifications in 16 neuron types. Bottom: Inclusion pattern of exon 11.5 from Tomioka et al. (2016), and agreement with our data. **D:** Raw data visualization of exon 11.5 alternative splicing in 5 neuron types. Top: Gene model. Bottom: Exonic and junction-spanning counts. Right: Exon 11.5 inclusion pattern from Tomioka et al. (2016) in the same neurons. **E:** Transcript quantification of daf-2 in the same neurons as C. The transcripts daf-2a, d, e, and f, (other) which do not include exon 11.5, were grouped together. Right: Exon 11.5 inclusion pattern from Tomioka et al. (2016).

To facilitate use by the scientific community, these data and analysis methods are available via a web portal at www.splicing.cengen.org. For raw data visualization, the user can select a gene or genomic region, and also choose the data to display: All individual samples are available, as well as the averaged data for each neuron type and the global data showing mean, maximum and minimum. For each data set displayed, the user can select whether to display the read counts, the exon-spanning reads, or both. In addition, due to our use of interoperable formats, our tracks can be imported to other genome browsers (such as WormBase or UCSC), and tracks generated by other projects can be displayed in our genome browser, allowing simultaneous examination of data from separate sources. For local quantification using MAJIQ, we display the results using VOILA. Finally, for transcript-level quantification, we developed a custom web application to display the results. For a single gene, the application displays both relative transcript usage and absolute transcript expression for all annotated transcripts. Multiple genes can be represented as a heatmap of transcript expression. These three tools offer users complementary levels of interpretation: quantification of transcript usage reflects the underlying biology of RNA processing. However, expression levels of complete transcripts are difficult to infer from short-read data and may be inaccurate (see Discussion). By contrast, local LSV-level quantification more directly reflects our measurements and is thus likely more accurate. However, for complex alternative transcripts, interpretation can be complicated by the need to consider multiple splice junctions simultaneously. Finally, the browser view does not offer rigorous quantification but can be used to examine the full context of a genomic region, including constitutive exons and non-coding RNAs.

### Axon guidance receptor gene *unc-40/DCC* is differentially spliced in specific neurons

Most gene models of splicing in *C. elegans* were obtained from sequencing bulk samples. However, if differential alternative splicing occurs in only a small number of cells, these rare splicing patterns might not be detected. To test this idea, we used manual inspection of raw data visualizations to examine well-studied neuronal genes.

We found that while a single transcript is annotated for the gene *unc-40*/DCC ^43^, our analysis detected two novel exons, exon 8.5 and exon 14.5 (Fig. 3A). In particular, exon 14.5 is preferentially included in AVM, whereas other neuron types (e.g. AVL, AWA) exclusively express the canonical splice variant, skipping exon 14.5 (Fig. 3B). Using RT-PCR, we validated the presence of the inclusion transcripts in cDNA extracted from whole animals, though at a lower level than the skipped transcript (Fig. 3C). Interestingly, the additional exons do not disrupt the open reading frame, and lead to insertions between known domains of the UNC-40 protein (Fig. 3D). To determine whether the inclusion of exon 14.5 represents a transcript unique to *C. elegans*, we examined the locus of the orthologs of *unc-40* in the closely related nematodes *Caenorhabditis briggsae* and *Caenorhabditis brenneri* (Fig. 3E), and examined bulk RNA-Seq data for those species available from Wormbase ^43^. We found that *C. briggsae* displays four annotated transcripts, with cassette exons corresponding to exons 8.5 and 14.5 of C. elegans. On the other hand, *C. brenneri* presents evidence of unannotated exons corresponding to exons 8.5 and 14.5. This finding suggests that other nematode species express a similar exon. The additional exons in Cbr-UNC-40 and Cbn-UNC-40 encode a protein sequences with high identity with exons 8.5 and 14.5 of Cel-UNC-40 (Fig. 3F). Thus, *C. elegans unc-40* has previously unannotated alternative transcripts, with conserved sequence and potential functional impact. In general, our analysis of splicing in individual neuron types can identify novel mRNA sequences.

**Figure 3:**
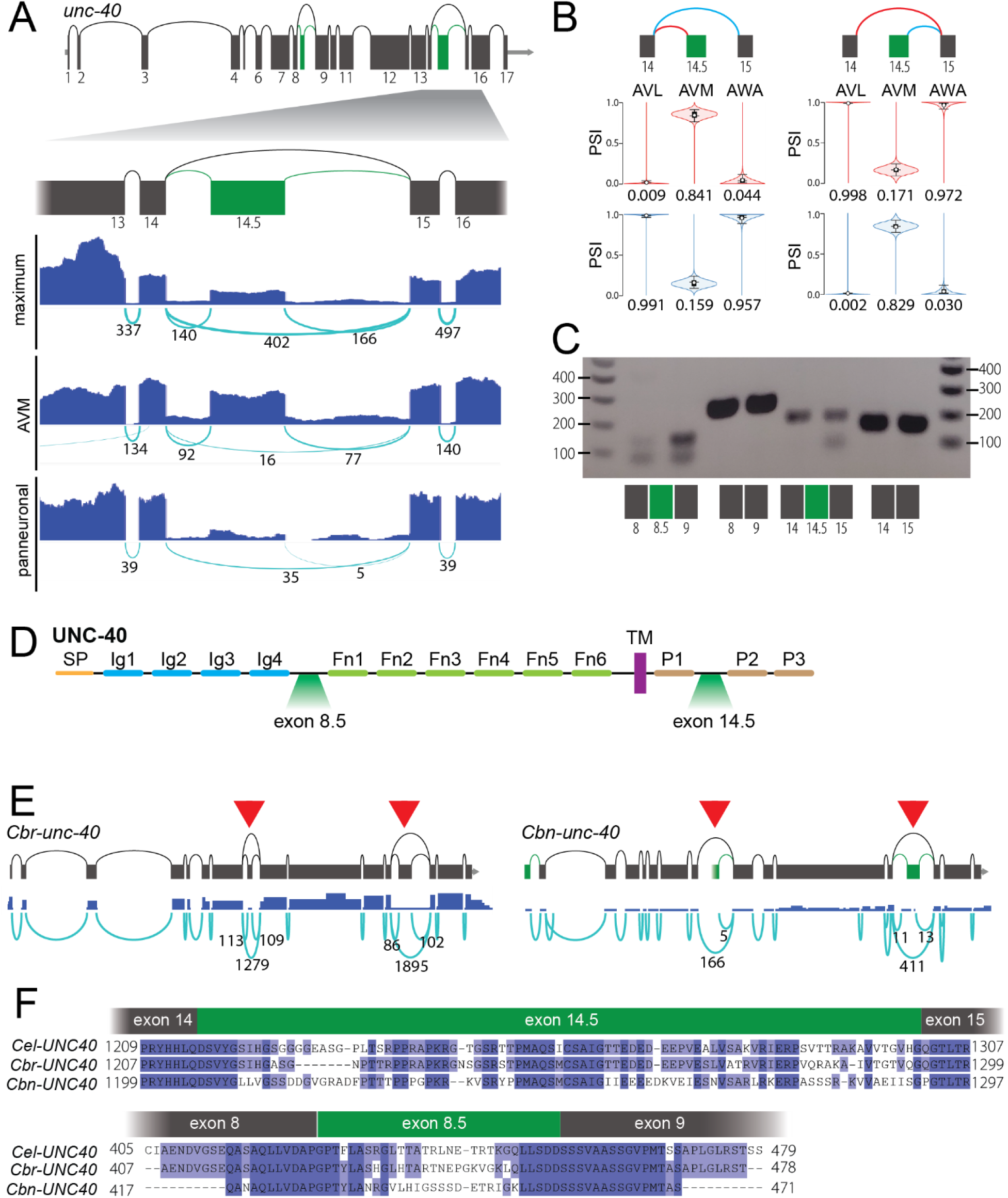
Detection of novel cassette exon in unc-40. **A:** Alternative expression of a novel cassette exon in unc-40. Top: Schematic of the gene structure, showing novel alternative exons (green) and the annotated constitutive exons (grey). Bottom: Genome browser representation near exon 14.5, in the maximum track, the AVM track, and in pan neuronal samples. **B:** Local quantification of exon 14.5 inclusion. **C:** Validation by RT-PCR of the novel exons in cDNA from whole animals. Exon-specific primers were used to amplify the indicated segments. Each pair represents independent amplifications. **D:** Protein structure of UNC-40, indicating the cassette exons vs known protein domains. SP: signal peptide, Ig: immunoglobulin domain, Fn: fibronectin domain, P: conserved intracellular motifs. **E:** Structure of the unc-40 orthologous genes in Caenorhabditis briggsae (Cbr-unc-40) and C. brenneri (Cbn-unc-40); red arrowheads denote orthologs of exon 8.5 and 14.5. Top: Annotated gene models (unannotated features in green). Bottom: RNA-Seq data. **F:** Alignment of the protein sequences of C. elegans UNC-40,C. briggsae UNC-40, and C. brenneri UNC-40 around exon 14.5 orthologs (top) and around exon 8.5 orthologs (bottom).

### *sax-3*/Robo and the homeobox factor *ceh-8* have novel alternative first exons

We also identified novel splice variants in the Slit receptor gene *sax-3*/ROBO. The *sax-3* annotation shows two transcripts, differing by 13 bp in exon 11 length (Fig. 4A) and 29 bp in the length of the annotated 5’ UTR (not shown in figure for clarity). We found a novel alternative splice site that shortens exon 9 by 15 bp. In addition, we detected a novel alternative first exon 5.5 (positioned between annotated exons 5 and 6). LSV quantification shows that the annotated alternative splice site in exon 11 is not used in the neurons sequenced here. By contrast, the novel alternative splice site in exon 9 and the alternative first exon 5.5 are both differentially expressed in broad subsets of neuron types in our data set (Fig. 4B, see AVL and AVM as examples). We confirmed the *in vivo* expression of both exon 9 splice sites and both alternative first exons by RT-PCR (Fig. 4C). Both of these novel events affect coding potential: The alternative splice site in exon 9 alters the amino acid sequence of the intracellular domain of SAX-3, whereas the alternative first exon 5.5 generates a short isoform (SAX-3S) lacking four of the five Ig domains, but encoding its own signal peptide in frame with the remainder of the protein (Fig. 4D). We examined the locus of the *C. briggsae* and *C. brenneri* orthologs, and find a remarkable conservation of the gene structure, including the novel exon 5.5 (Fig. 4E). These orthologous exons 5.5 encode an identical amino acid sequence (Fig. 4F).

**Figure 4:**
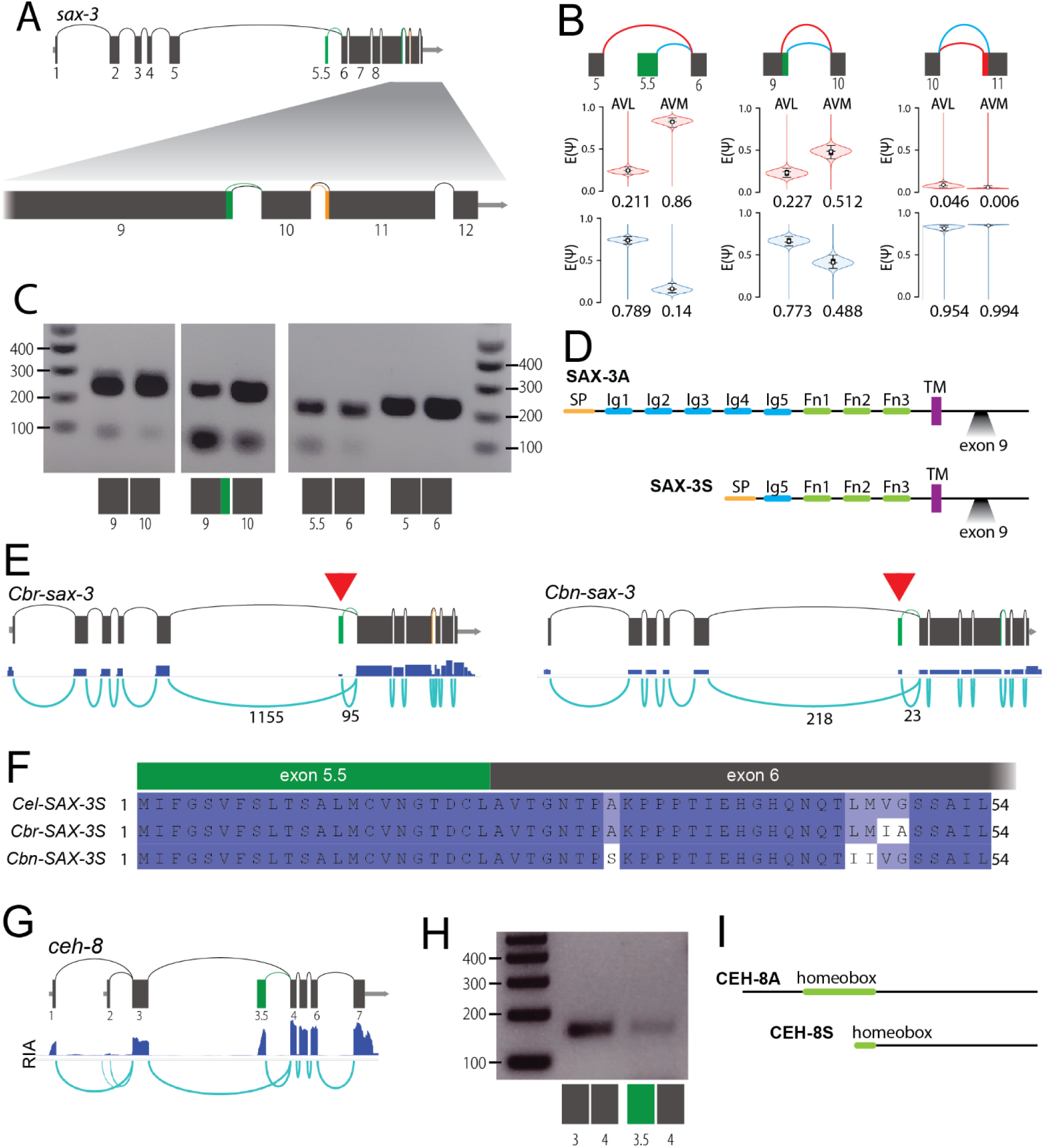
Detection of novel alternative first exons in sax-3 and ceh-8. **A:** Structure of the gene sax-3/ROBO. The novel exon 5.5 and the novel alternative splice site at the end of exon 9 are represented in green, the annotated alternative splice site at the start of exon 11 is represented in orange. Bottom: Enlarged 3’ end of the locus. **B:** Local quantification of LSVs for the novel alternative first exon 5.5 (left), the novel alternative 5’ splice site at the end of exon 9 (middle), and the annotated alternative 3’ splice site at the start of exon 11 (right). Quantifications are displayed for the representative neurons AVL and AVM. **C:** Validation by RT-PCR of the novel exons in cDNA from whole animals. Exon-specific primers were used to amplify the indicated segments. Each pair represents independent amplifications. **D:** Protein structure of SAX-3, indicating the impact of the novel splice variants. SP: signal peptide, Ig: immunoglobulin domain, Fn: fibronectin domain. Top: Overall structure of annotated isoform SAX-3A, showing position of novel splice site in exon 9. Bottom: Overall structure of the novel short isoform starting at novel exon 5.5 (SAX-3S), showing position of novel splice site in exon 9. **E:** Structure of the orthologs of sax-3 inc C. briggsae (Cbr-sax-3) and C. brenneri (Cbn-sax-3). Top: Gene models. Bottom: RNA-Seq data. Red arrowheads denote the orthologs of exon 5.5. **F:** Protein sequence alignment of the short isoforms of SAX-3 orthologs from C. elegans, C. briggsae, and C. brenneri. **G:** Alternative expression of the novel first exon 3.5 in ceh-8. Raw data visualization in neuron RIA, with novel alternative first exon 3.5 (green) vs annotated transcript (grey). **H:** Validation by RT-PCR of the novel exons in cDNA from whole animals. Exon-specific primers were used to amplify the indicated segments. Each pair represents independent amplifications. **I:** Structure of the annotated protein isoform CEH-8A and the novel isoform starting at N-terminal alternative exon 3.5 CEH-8S.

Finally, we identified a novel alternative first exon in the homeobox transcription factor *ceh-8* (Fig. 4G, H). Interestingly, this transcript would lead to translation of a protein with a truncated homeobox domain (Fig. 4I). The *ceh-8* locus is not well conserved in *C. briggsae* and *C. brenneri*, precluding direct comparison of alternative transcripts. Together, these examples demonstrate that our approach can detect novel alternative first exons. Such events also contribute to transcript diversity, but are generated by different biological mechanisms than other forms of alternative splicing. While most alternative splicing is performed by the spliceosome, alternative first exons are the result of alternative promoter usage.

### Global detection of novel splicing events across neuron types

Given these examples of novel isoform detection, we sought to identify candidate novel splice junctions across our data set. We used STAR to generate a preliminary list of 1,026,619 unannotated splice junctions ^50^. We filtered these junctions using multiple criteria to focus on well-supported novel junctions, for example by requiring high expression relative to neighbor genes (see Methods). We also leveraged the biological replicates in our data set, requiring novel splice junctions to be present in at least half the samples from a single neuron type, minimum of two. This analysis yielded 1,722 novel junctions (Table S2). Attaching novel junctions to annotated genes is a transcript discovery task which is challenging using short reads, and current tools do not reach high accuracy ^47^. However, each novel junction must belong to a gene in its immediate neighborhood; we provide a list of all neighbor genes (Table S2). We also sought to estimate the number of genes containing novel splicing events, without precise knowledge of which gene each junction belongs to. As multiple events may belong to the same novel transcript, we expect the number of affected genes to be less than the number of novel splice junctions. For each junction, we focused on the set of neighbor genes and randomly selected one gene per junction neighborhood. We repeated the procedure 1,000 times and found an average of 1,361 genes containing at least one novel junction. Overall, this analysis indicates that many novel splice sites and mRNA isoforms remain to be described, and provides a candidate list for future study.

### Detection of differential AS between neuron types identifies genes associated with neuronal excitability

Our analysis of known differential alternative splicing events (*ric-4, elp-1, daf-2*; Figs. 1,2) and identification of novel events (*unc-40, sax-3, ceh-8*; Figs. 3,4) demonstrate that our data set can be used to identify instances of differential alternative splicing (DAS) in the *C. elegans* nervous system. To identify candidate DAS events across all genes and neuron types, we quantified differential event usage using MAJIQ ^46^. Specifically, within its Local Splicing Variation framework, MAJIQ models the relative usage of splice junctions that share a common source or target exon. Across the 55 neuron types in our data set, we detected 1,940 genes displaying DAS. To validate these findings, we compared the genes detected here to a list of 542 genes compiled from the literature (Table S3, see Methods) and found a large overlap of 461 genes (85%; Fig. 5A). This comparison indicates that the novel instances of DAS we detect are indeed strong candidates.

**Figure 5:**
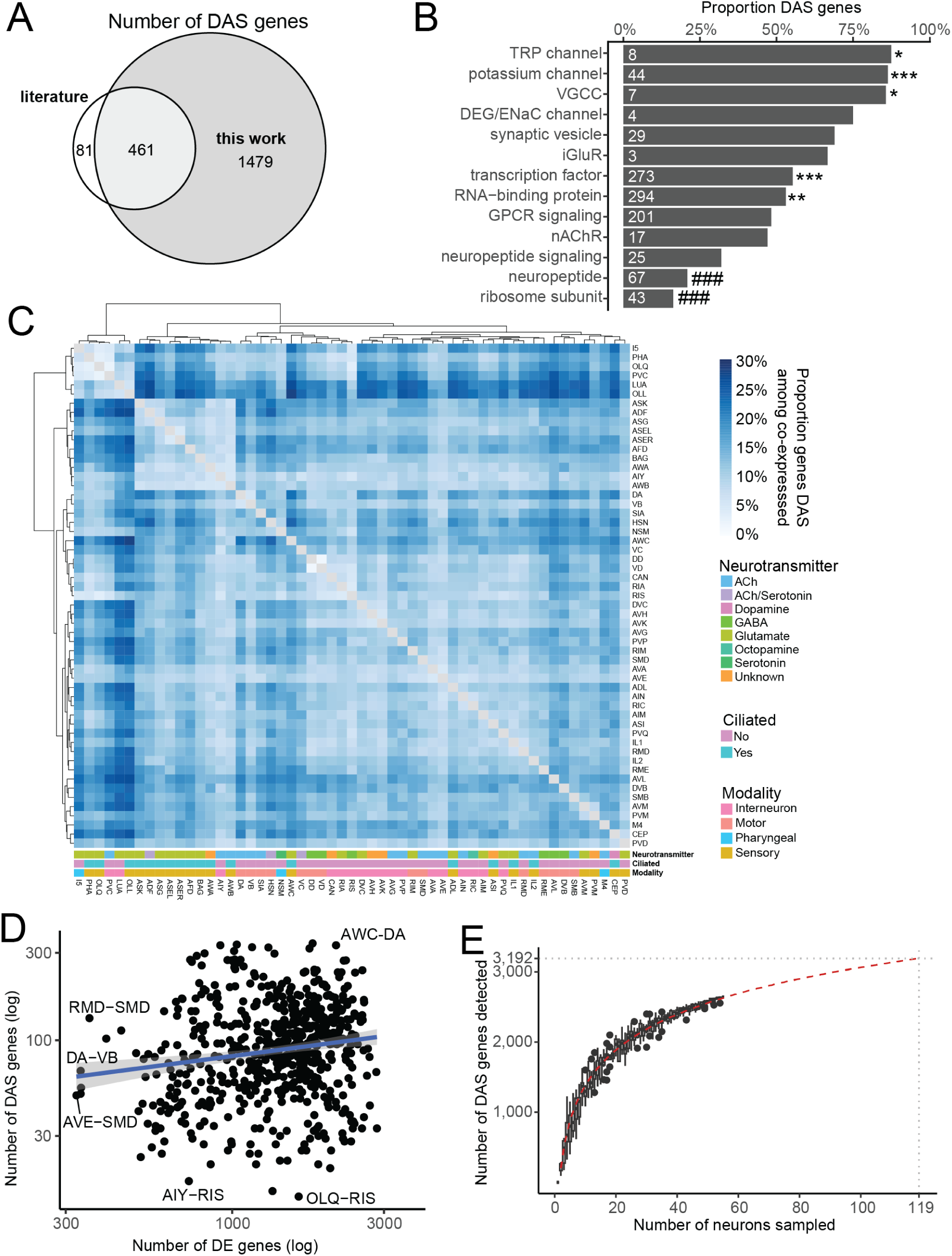
Differential AS between neuron types. **A:** Euler plot of the number of genes presenting differential alternative splicing (DAS) in this analysis, and from a literature review (Table S2). **B:** Proportion of genes DAS within families of neuronally significant genes 52. The number of genes in the family is indicated inside each bar. Hypergeometric test: * Significantly enriched, # significantly depleted. **C:** Proportion of DAS genes per neuron pair, among genes co-expressed in the neurons of this pair. **D:** Number of DAS genes plotted against number of differentially expressed genes for each neuron pair. **E:** Estimate of the number of genes in which DAS can be detected relative to the number of neuron types sequenced.

Next, we sought to explore the function of candidate DAS genes. Gene Ontology analysis ^51^ showed enrichment of terms related to neuronal function (Fig. S1A). To investigate the specific role of DAS, we examined major neuronal functional gene classes ^52^. We found that the prevalence of DAS is highly variable by gene class. For example, the majority of potassium channels and voltage-gated calcium channels undergo differential alternative splicing, whereas ribosome subunits and neuropeptide-encoding genes tend to be similarly spliced across neuron types (Fig. 5B). This analysis suggests that DAS is preferentially used to fine tune neuronal excitability, enhancing functional diversity among different neuronal types.

### Global patterns and prevalence of DAS

With a list of DAS events in hand (Fig. 5A), we examined global patterns of DAS across neuron types. We performed pairwise comparisons of all neuron types in our data set, and assessed the proportion of DAS genes among the genes co-expressed in both neurons of the pair (Fig. 5C). Clustering this data revealed that that a group of 10 ciliated sensory neurons (ASK, ADF, ASG, ASEL, ASER, AFD, BAG, AWA, AIY, AWB) have similar global patterns of AS. In addition, some pairs of neurons with similar functions have similar splicing profiles (DA-VB, DD-VD, AVM-PVM, AVH-AVK) (Fig. S1B). Interestingly, another group of 6 neurons comprising I5, LUA, OLQ, OLL, PHA, and PVC also display similar AS profiles (Fig. 5C), even though they do not share known functional or morphological characteristics. These data suggest that at least in some cases, neurons with similar characteristics adopt similar patterns of DAS, presumably to support specific functional specialization.

Besides DAS, gene expression patterns are highly correlated with neuron type. In fact, gene expression patterns alone can be used to group single cells into clusters corresponding to individual neuron types ^32^. Given the very strong association between gene expression and neuron type, we wondered if DAS patterns and gene expression patterns are related. In this case, both gene expression and alternative splicing might encode the same information reflecting the underlying structure of neuronal cell types. To test this model, we compared (for each neuron pair) the most strongly differentially expressed (DE) genes to the most strongly DAS genes, and found limited overlap (Fig. S1C). In addition, for each pair of neurons, we compared the number of DE genes to the number of DAS genes and found only a weak correlation (Fig. 5D). Together, this analysis indicates that DAS and gene expression are two largely independent dimensions of neuron type identity ^53^.

We then aimed to determine the number of DAS genes in the entire nervous system. As our dataset covers 55 neuron types out of 119 classes (118 canonical classes, plus ASEL/ASER), we used a downsampling approach to estimate the number of detected DAS genes depending on the number of neuron types sequenced (Fig. 5E). By projection, we estimate that 3,192 genes are differentially alternatively spliced within the nervous system (see Methods), corresponding to about one quarter of all genes expressed in neurons.

### RNA features of alternative splicing

Alternative splicing can take many forms, with implications for both regulatory mechanisms and biological effects. To assess the representation of different alternative splice types in the *C. elegans* nervous system, we grouped events into canonical event types (Fig. 6A). We found that alternative splice sites, cassette exons, and alternative first exons are well represented in the genome. By contrast, alternative last exons, intron retention, and coordinated multiple exon skipping are relatively rare. We assessed the prevalence of differential alternative splicing within each event type and found that DAS is well-represented among all forms of alternative splicing. This analysis indicates that neurons use all available mechanisms to increase their molecular diversity.

**Figure 6:**
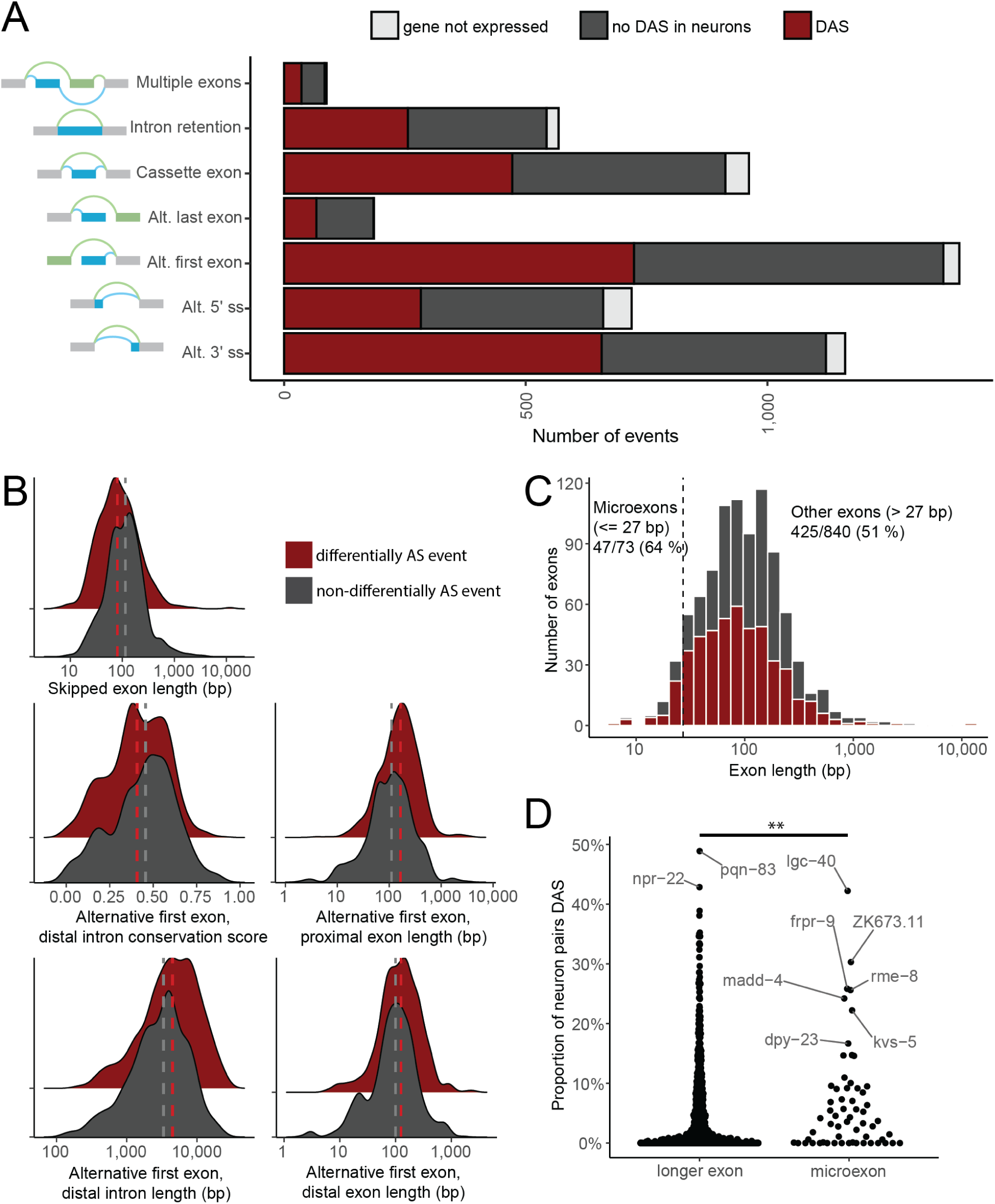
Landscape of alternative splicing event types. **A:** Number of events by types. Bar length indicates the number of canonical events of each type predicted from genome annotation. Dark grey indicates the events that did not display DAS between neuron types in our dataset; dark red indicates events with DAS in at least one pair of neurons; light grey indicates events in genes that could not be measured in our neuronal dataset. **B:** Distribution (density plot) of GC content, conservation score, and length in splicing events. DAS events (red) vs and non-DAS events (grey). Features with a statistically significant difference between DAS and non-DAS events are represented here. The vertical dashed lines represent the median of each group. **C:** Histogram of cassette exon lengths, separating DAS exons (red) and non-DAS exons (grey). The vertical dashed bar delimits microexons. **D:** Each dot represents one exon. The vertical axis shows the proportion of neuron pairs where the exon is differentially AS between the two neuron types (proportion calculated among the neuron pairs in which the exon-containing gene is expressed in both neurons of the pair). ** p < 0.01 (Wilcoxon test).

Previous work has found a role of sequence features in distinguishing AS events from constitutive splicing ^13, 14, 15, 16, 17^. Do sequence features also affect *differential* AS? To address this question, we examined the association of broad sequence features with differential AS. For each event type, we delineated the genomic regions composing the event locus. For example, for cassette exons, we considered the alternative exon as well as the two flanking introns. For each of these genomic regions, we measured its length, GC content, and conservation score. We compared the resulting values between differentially spliced events vs. events where we did not detect differential AS. In total, this resulted in 72 comparisons (Table S4), of which 5 were statistically significant (Fig. 6B, Table 1). Strikingly, four of the significant differences were for alternative first exons. Alternative first exons that display differential AS between neuron types have longer introns, a less conserved distal intron, and a longer distal exon. Alternative first exons are features of transcriptional regulation rather than post-transcriptional splicing. These data indicate that sequence features surrounding first exons may be highly variable to support fine-tuning expression of alternative isoforms at the level of transcription.

**Table 1:** Sequence features presenting a statistically significant difference between dAS events and non-dAS events.

| Event type | Feature | Median when dAS | Median when not dAS | p-value |  |
| --- | --- | --- | --- | --- | --- |
| Alt. first exon | Distal exon length | 124 bp | 100 bp | 1.20E-05 | *** |
| Alt. first exon | Distal intron length | 4,468 bp | 3,340 bp | 4.80E-12 | *** |
| Alt. first exon | Distal intron conservation | 0.407 | 0.456 | 0.035 | * |
| Alt. first exon | Proximal intron length | 166 bp | 112 bp | 8.30E-13 | *** |
| Cassette exon | Exon length | 80 bp | 114 bp | 7.00E-05 | *** |

By contrast, cassette exons presenting differential AS in the nervous system appeared shorter on average. This observation is reminiscent of recent findings that microexons, with a length of 27 bp or less, are differentially spliced between *C. elegans* tissues ^37^ and frequently display neuron-specific inclusion ^54^. We thus focused more closely on the differential splicing of microexons and found that, of the 73 microexons measurable in our dataset, 47 (64%) displayed differential splicing in the nervous system, as opposed to 51% of the larger alternative exons (Fig. 6C) (p = 0.032). Furthermore, we asked if microexons showed more differential AS than other exons (Fig. 6D). Indeed, microexons were on average DAS in 6.6% of the neuron pairs tested, as opposed to 4.7% for longer exons (p = 0.0047).

Overall, our analysis indicates that differential alternative splicing is associated with specific features of the pre-mRNA. For alternative first exons, which are associated with an increase in intron and exon length, these features likely include transcriptional start sites. Cassette exons and microexons displaying DAS, by contrast, tend to be shorter than constitutive exons. In this case, shortness may be a result of selection for protein integrity, with only minor insertions or deletions well-tolerated.

### Splicing Regulatory Network

Differential alternative splicing between neuron types is likely regulated in part by differential expression of splice factor (SF) genes. Our data enables the concurrent measurement of DAS and of gene expression in the same samples across 55 neuron types. We reasoned that these measurements might enable elucidation of a splicing regulatory network that links splice factor expression to DAS. To perform this analysis, we first sought a single measurement that could quantify DAS across genes and neuron types. For this purpose, we restricted our analysis to cassette exons—exons which are either included or excluded from the final transcript (e.g., *unc-40* cassette exon in Fig. 3). For each cassette exon, we computed a Percent Spliced-In measure (PSI) which captures the extent to which the exon is included in each individual neuron type (see Methods). We also compiled a list of putative SF genes (Table S5) and quantified their expression. Our list includes well-studied splice factors, as well as genes with only a speculative role in splicing regulation. Since SF genes themselves are heavily AS, we separately quantified expression of each SF transcript. These two measurements—PSI and SF expression—constitute the input data for our model.

We constructed a covariance matrix which assesses, for each cassette exon PSI and each SF transcript, how they covary over all 55 neuronal types in our data. In principle, the inverse of this covariance matrix is a precision matrix corresponding to the splicing regulatory network. However, the covariance matrix cannot be directly inverted due to the underdetermined system of equations that comprises many more covariates (SFs and PSIs) than observations (neuron samples). Thus, we sought to estimate the precision matrix (Fig. 7A).

**Figure 7:**
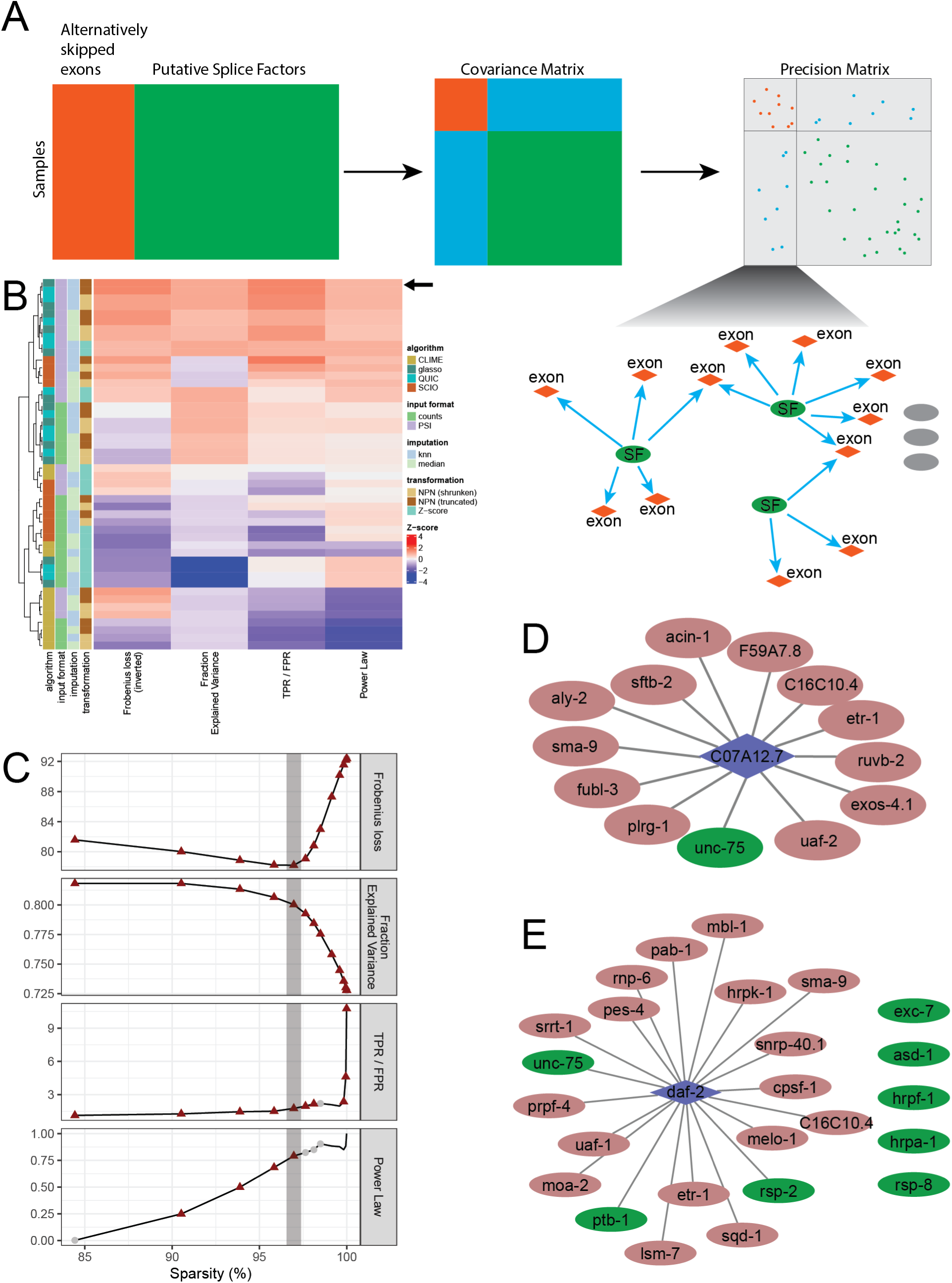
Splicing Regulatory Network. **A:** Schematic representation of the approach. **B:** Comparison of metrics for selected parameters. The four metrics are represented across all methods. The Frobenius loss was inverted such that for all metrics, higher is better. The arrow represents the selected “best” method. **C:** The four metrics plotted against sparsity for a range of penalties. The grey line highlights the selected penalty of 0.25, corresponding to a sparsity of 97%. The red triangles correspond to values significantly different from the permuted data. **D, E:** Subnetworks centered on exon 5 of C07A12.7 (D) and exon 11.5 of daf-2 (E). Blue diamond: cassette exon, red ellipse: putative splice factor displaying a non-zero weight in the network, green ellipse: splice factor identified from the literature to regulate this exon.

To select our method of precision matrix estimation, and optimize the hyperparameters, we used 5-fold cross-validation and computed four metrics (Fig. S2A, see Methods). First, the Frobenius norm loss reflects the ability of the method to capture correlations. Second, the Fraction Explained Variance (FEV) of the PSI reconstruction using regression coefficients ^55^ reflects the ability to capture relationships between SF expression and PSI. Third, we compiled a ground truth dataset of splicing regulatory interactions for comparison to the network structure. Finally, we used a scale-free criterion on the structure of the network ^56^. Using these metrics, we compared the following combinations of approaches. For precision matrix estimation, we considered the glasso ^57^, QUIC ^58^, CLIME ^59^, and SCIO ^60^ algorithms. For input to these algorithms, we considered either the measured PSI or reconstructed counts. For normalizing the range of these inputs, we considered applying a Z-score or nonparanormal transformation ^61^. Lastly, to handle missing values we considered k-nearest neighbors interpolation or median interpolation. Based on our evaluation metrics in a cross-validation, we chose to use the following methods: glasso as the precision matrix estimator algorithm, PSI as input, nonparanormal truncated transformation, and k-nearest neighbors imputation (Fig. 7B, arrow), selecting 4-nearest neighbors (Fig. S2B).

We then used all the training and validation data, together with the selected methods, to train a final model. We selected an optimal penalty for the glasso algorithm and assessed the model’s performance on the as yet unseen test data using a permutation test approach. First, we permuted the model by randomizing the input cassette exon quantifications between samples in the training and validation data. For each permutated dataset, we tested a range of glasso penalty values. We then recalculated the model based on these unstructured data. We compared the performance metrics of these randomized models to that of the model trained on the unaltered data, at each glasso penalty value. For most metrics, the model fails to appropriately reconstruct the validation set when the input data is unstructured, indicating that the model captures relevant relationships (Fig. S2C). We also calculated a permutation test p-value that reflects the relevance of the model for each metric and selected a glasso penalty value that optimized performance (Fig. 7C). Our final model establishes a virtual network relating putative splice factors to splicing of cassette exons.

One application of our splicing network is discovery of potential splice factors. To this end, we examined the putative splice factors with the most central position in the network, ordered by number of connections to cassette exons. We found that many of the most central genes are indeed splice factors known to act in *C. elegans* neurons (Table 2). For example, *mbl-1*/Muscleblind (Norris et al. 2017), the hnRNP *hrpa-1* (Tomioka et al. 2016), and the CELF family gene *unc-75* (Kuroyanagi et al. 2013a) are all known splice factors in *C. elegans* neurons. Our analysis indicates that these key factors likely regulate differential alternative splicing across genes and neuron types, at least of cassette exons. Other central nodes are genes not previously known to function in neuronal splicing. For example, we identified the CELF family gene *etr-1* as a key splice factor. *etr-1* has known roles as a splice factor in muscle (Milne and Hodgkin 1999; Ochs et al. 2022), and our recent data from single-cell RNA-Seq suggests expression in a restricted set of neurons (Taylor et al. 2021). Together, these data suggest that *etr-1* might play a novel role in regulating neuronal DAS, similar to its known role in muscle. Our analysis also placed the transcription factor *sma-9* and *melo-1*/periphilin 1 in a central position. Neither *sma-9* nor *melo-1* has an established role in splicing regulation. Thus, central nodes in our network, besides known splice factors, may indicate novel regulators of alternative splicing in neurons.

**Table 2:**
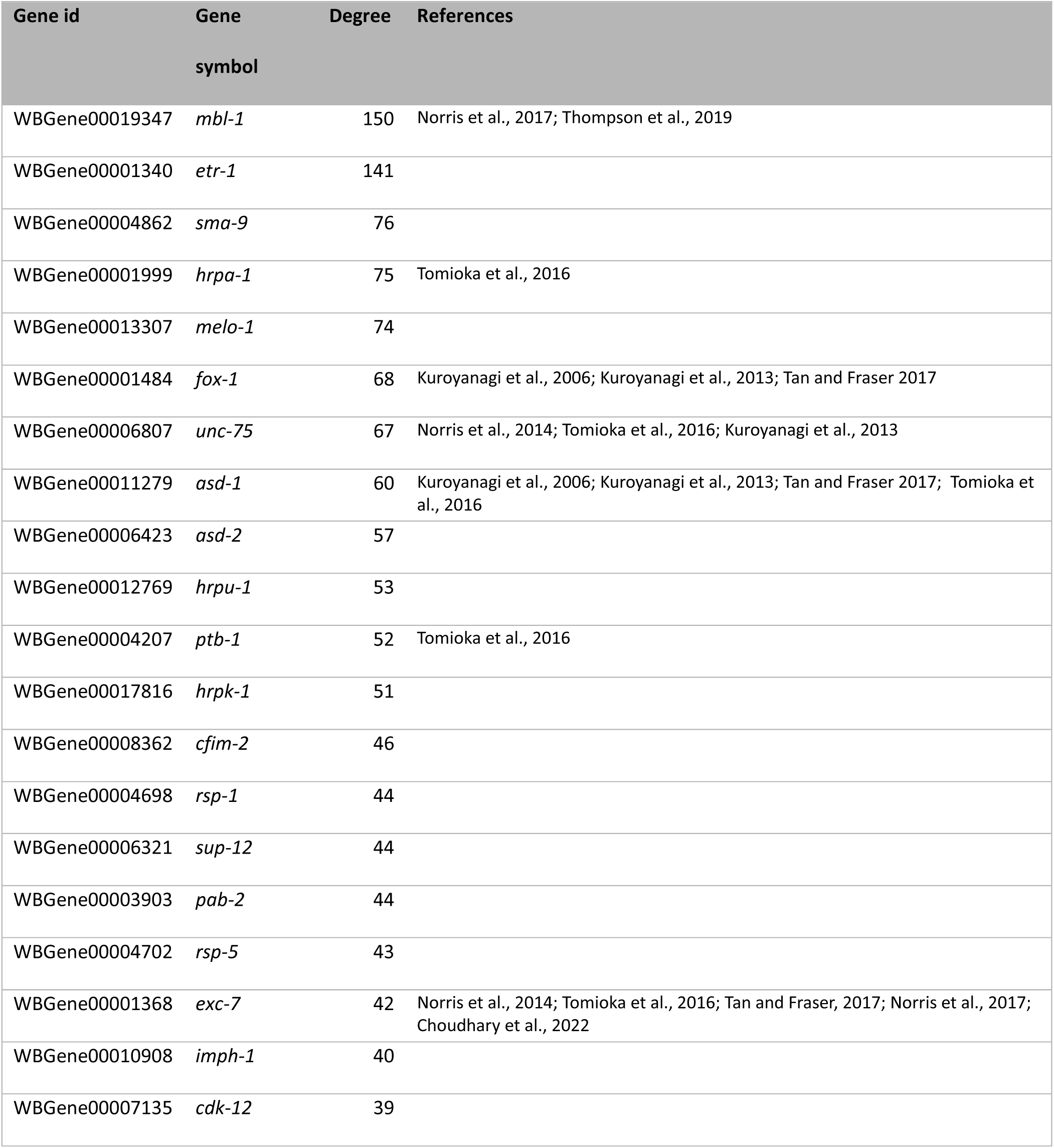
Top 20 putative splice factors with highest network degree.

**Table 3:**
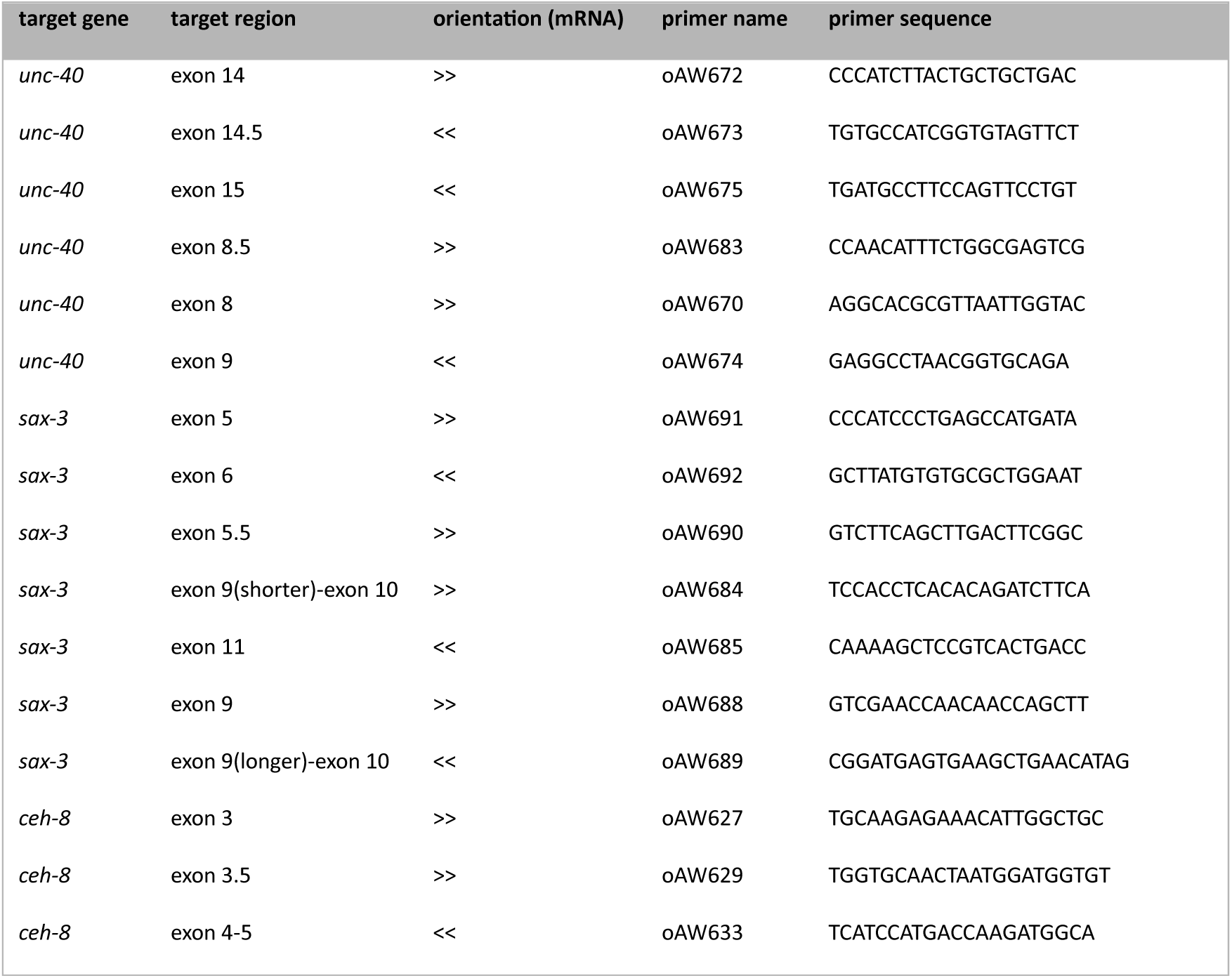
Primers used for RT-PCR.

A second application of our network is to analyze the regulation of specific cassette exons. We examined the subnetwork of putative SFs connected to *C07A12.7* exon 5 (Fig. 7D) and *daf-2* exon 11.5 (Fig. 7E). For *C07A12.7*, the known regulation by *unc-75* (Kuroyanagi et al. 2013b) was correctly detected. In addition, 12 other putative interactors appeared in the network: *acin-1*, *aly-2*, *C16C10.4*, *etr-1*, *exos-4.1*, *F59A7.8*, *fubl-3*, *plrg-1*, *ruvb-2*, *sftb-2*, *sma-9*, and *uaf-2*. Similarly, for *daf-2* exon 11.5, we detected the known regulation by *ptb-1*, *rsp-2*, and *unc-75* (we did not detect regulation by *asd-1*, *exc-7*, *hrpa-1*, *hrpf-1*, and *rsp-8*) ^49, 62^. We obtained an additional 17 predicted interactors: *C16C10.4*, *cpsf-1*, *etr-1*, *hrpk-1*, *lsm-7*, *mbl-1*, *melo-1*, *moa-2*, *pab-1*, *pes-4*, *prpf-4*, *rnp-6*, *sma-9*, *snrp-40.1*, *sqd-1*, *srrt-1*, and *uaf-1*. In addition, we examined the subnetwork of putative splice factors connected to *unc-16*. We quantified 6 separate events, corresponding to 4 exons (two pairs of events correspond to the same exon with differing flanking introns) in the gene *unc-16*, represented as individual nodes in the subnetwork (Fig. S2D). We identified the known regulation by *unc-75* and *exc-7*, but did not detect regulation by *prp-40* ^63, 64, 65^. In addition, we found 58 network connections that do not correspond to known regulatory interactions. Thus, our splicing network can generate detailed hypotheses about the regulation of specific splicing events.

## Discussion

In this study, we present an atlas of alternative splicing (AS) in the *C. elegans* nervous system, at single neuron type resolution, across 55 neuron types. We develop a toolset of analytic approaches and a website to facilitate their use by the research community. We show that our approach yields results in agreement with existing data, and identifies multiple new examples of alternative splicing in genes with key roles in neuronal function. Systematic quantification reveals broad patterns of differential AS, particularly affecting ion channels, and helping to shape the landscape of neuronal mRNAs separately from gene expression. We provide a broad description of the *C. elegans* neuronal alternative transcriptome, and observe that microexons are notably differentially alternatively spliced (DAS) in the nervous system. Finally, we compute a splicing regulatory network to formulate new hypotheses on splice factor regulation of differential alternative splicing, focused on cassette exons.

With the wider availability of single-cell RNA-Seq methods, large efforts have been made in the recent years to establish atlases of gene expression in *C. elegans* ^29, 30, 31, 32, 33, 34, 35^. However, these sequencing approaches often present a strong 3’ end bias which make them unsuitable for AS analysis. A more restricted set of studies have focused on AS. In *C. elegans*, these studies have been performed mostly at the level of whole animals, or at the level of tissues ^37, 38, 39^. Studies in other organisms have examined AS at the tissue ^19^ or single-cell level in the nervous system ^3, 66, 67, 68, 69, 70^. A feature of our dataset is its genome-wide scope and its high resolution analysis of AS across many individual neuronal types. Thus, this analysis complements and extends previous studies.

Our analysis is broadly consistent with previous work, while identifying thousands of new features. Two apparent discrepancies are our global counts of intron retention and alternative first exons. For intron retention, we found relatively fewer instances than a parallel analysis of the same data (ref. Wolfe et al. manuscript). In this case, our use of the tool SUPPA only measures intron retention relative to annotated transcripts, and does not detect unannotated instances. For alternative first exons, we found more instances than an analysis of a separate data set ^37^. In particular, our use of the tool SUPPA considers multiple alternative first exons in the same gene as distinct events, rather than grouping them as a single event. Overall, these discrepancies highlight the difficulty of computationally interpret alternative splicing at genome scale. For this reason, we encourage inspection of the raw data visualization we provide to guide investigation of splicing at any specific locus.

We used a computational approach to define a Splicing Regulatory network. A challenge in evaluating the performance of such an approach is the availability of a ground truth. Although we used a compiled list of interactions between splice factors and splice events as one metric, some of these data are not compiled from neurons, and even neuronal data are not at the resolution of individual neuron types, making it difficult to assess our model using that approach alone. Instead, we adopted an innovative approach to select the hyperparameters of the model using four separate criteria. In contrast, previous attempts did not compare alternative approaches, and either did not justify the initial selection of hyperparameters ^19, 22^, or used a single criterion ^3, 20^. Our principled tuning method may be useful for other models with limited access to ground truth. Overall, our final network captures many known interactions between putative splice factors and splice events, and constitutes a powerful hypothesis generation tool for discovery of novel splicing regulatory mechanisms.

Most previous effort to model the role of splice factors on DAS were performed with RNA-Seq data set either derived from whole animals ^22^ or from tissue-specific samples ^19, 20, 24, 27, 71, 72^. The recent progress in single-cell RNA-Seq has facilitated higher resolution studies ^3, 18^. In particular, current progress in single-cell full length sequencing offers great opportunities to extend this analysis ^73, 74, 75^. These single cell approaches allow for more specificity and potentially allow for whole-transcript analysis, enabling higher-quality atlases, and deeper analysis of splicing networks, in the near future.

Our approach has cataloged DAS in single neuron types for almost half the canonical neuron types in the *C. elegans* nervous system. Our data indicate that alternative splicing affects the function of key neuronal genes, and reveals substantial novel splicing diversity. These splicing events might control subtle, cell-specific alterations of neuronal form and function that are not accessible by broader forms of genome regulation. Thus, we expect studies of gene function to be informed by these data about differential alternative splicing in specific neuron types. In addition, from the perspective of splicing itself, we use the diversity we have discovered to model regulatory mechanisms that mediate the control of differential alternative splicing.

## Methods

### FACS isolation and sequencing

For single-cell type bulk RNA-Seq, *C. elegans* strains expressing a fluorescent protein or combination of fluorescent proteins in a single neuron type were dissociated and sorted into Trizol as described previously ^32, 40^. RNA was extracted and sequencing libraries were prepared using the Ovation® SoLo® RNA-Seq library preparation kit, yielding even coverage along the gene body, as described previously ^41^. Libraries were sequenced with an Illumina HiSeq 2500 or NovaSeq 6000 (Table S1). The dataset covers 211 samples corresponding to 55 neuron types, and an additional 8 control samples from pan-neuronal sorts. All neuron types were sequenced in 3-6 replicates, except ADF, M4 (1 replicate each), OLL and PVD (2 replicates). Four samples failed quality control and were excluded from subsequent analyses. Following trimming, the RNA-Seq reads were aligned to the *C. elegans* genome (Wormbase WS289) using STAR 2.7.7a ^50^ with option (all other settings left to `--outFilterMatchNminOverLread 0.3` default). Deduplication was performed using UMI-tools ^76^. The pipeline code is available at: https://github.com/cengenproject/bulk_align.

### RT-PCR

For RT-PCR, mixed stage N2 *C. elegans* were grown following standard methods ^77^. Plates were washed with M9 buffer, and 100 µL of worm suspension was added to 400 µL of Trizol and immediately frozen in liquid nitrogen. Samples were stored at −80°C. RNA extraction was performed with chloroform in Phase Lock Gel Heavy tubes (QuantaBio), treated with DNase I (Thermo Fisher) and purified with the Macherey-Nagel Nucleospin kit following manufacturer’s instructions. Finally, cDNA was synthetized using the Affinity Script Multiple Temperature cDNA Synthesis kit (Agilent), following manufacturer’s instructions with oligo-dT primers. The resulting cDNA was stored at −20°C. RT-PCR reactions were performed with Taq polymerase (New England Biolabs) with cDNA diluted to 10 ng/µL, following the manufacturer’s protocol.

### Discovery of novel splice junctions

STAR produces junction files, providing a list of splice junctions detected in the processed sample, along with the measured count, annotation status, and other information ^50^. We only considered novel junctions (not present in the annotation), that were flanked by canonical splice site motifs. In addition, we only considered splice junctions supported by reads with 12 bp overhang on each side of the junction (STAR’s default value for `–-outSJfilterOverhangMin`). We defined the neighborhood of a splice junction as the set of genes within 60 bp of either splice site, regardless of the strand. We then filtered the junctions in each sample fulfilling the following criteria:

- Junction no longer than 1,000 bp
- At least 2 supporting reads (uniquely mapped, with at least 12 bp overhang following STAR default) supporting that junction
- Not in the neighborhood of an rRNA gene
- In the neighborhood of a protein-coding gene, long-non-coding RNA, or pseudogene
- Has at least 20 % as many reads as the most highly detected splice junction from the neighbor genes

After processing each sample with the above filters, we aggregated the junctions across samples and conducted a second round of filtering. We kept novel splice junctions that were detected in at least half the samples from a single neuron type (with a minimum of two samples from a single neuron type). This analysis identified 1,722 novel junctions robustly expressed in our dataset. Reliably attributing each junction to a novel transcript is challenging with available methods^47^. Instead, we only attempted to estimate the total number of genes containing novel junctions, without determining their precise identity. To this end, we determined the set of genes neighboring each novel junction, randomly sampled a single one, and evaluated the number of genes with novel junctions. By repeating this procedure 1,000 times, we estimated the total number of genes expressing novel junctions. The corresponding code is available at: https://github.com/cengenproject/novel_junctions.

### dAS with MAJIQ

Local quantification of AS and the analysis of differential AS were performed with MAJIQ 2.3 ^46^. We built a configuration file using the reference annotation from Wormbase (WS289), strandness forward, and one experiment per neuron type (grouping the biological replicates by experiment). We subsequently ran `majiq build` with parameter `--min-experiments 2` (i.e. a splice junction is retained if it passes filters in at least two replicates from the same neuron), keeping the other options to their default values. We performed PSI quantification, and delta PSI quantification between each pair of neuron types with the default parameters.

For the subsequent analysis, we grouped the resulting delta PSI files from all neuron pairs, obtaining 16,379,082 individual comparisons (for a given LSV in a given pair of neurons). We filtered comparisons to retain only those where the LSV-containing gene was expressed in both neurons of the pair, using the threshold “3” we previously defined based on single-cell RNA-Seq ^32^, resulting in 8,783,582 comparisons (corresponding to 3,787 measurable genes). We define a comparison as DAS if the “probability of not changing” (computed by MAJIQ deltapsi) is lower than 0.05, and the “probability of changing” is higher than 0.5, corresponding to 928,985 comparisons. Finally, we define a gene as DAS if it contains at least one DAS comparison, yielding 1,940 DAS genes.

To predict genes expected to display DAS in the nervous system, we compiled a list of 11 studies reporting individual genes ^78, 79, 80, 81, 82, 83, 84^ or performing transcriptomic analyses in mutant backgrounds disrupting AS in neurons ^64, 85, 86, 87^. This resulted in a list of 759 genes (Table S3), of which 542 are measurable in our dataset (expressed in the neurons sequenced and quantified by MAJIQ). Gene Ontology analysis was performed using a background list of 10,312 genes expressed in at least two neurons sequenced here (as per the threshold above).

To compare the differentially AS to differentially expressed (DE) genes, the DE genes were obtained from integration of single-cell RNA-Seq data with this dataset, as described in ^40^. For each neuron pair, the DE genes were ordered by absolute fold change and the genes with the 100 highest values were selected. The DAS genes were ordered by absolute delta PSI of their highest LSV, and the 100 genes containing the highest values were selected. Out of 595 neuron pairs, we could select a top 100 DAS genes for 432 pairs; 15 had 101 genes (because of a tie in the highest delta PSI), 148 pairs did not have 100 DAS genes. We only represented pairs where we could select 100 top DAS genes.

To predict the total number of DAS genes in the nervous system, we randomly selected between 2 and 55 neuron types among those sequenced, and estimated the number of DAS genes that could be detected. We repeated the procedure 10 times for each number of neurons. We then performed a linear regression of the number of DAS genes detected vs the logarithm-transformed number of neurons subsampled, yielding the relationship *N_genes_* = −302 + 731 ⋅ log (*N_neurons_*). The total number of DAS genes for 119 neuron types is then 3192. We applied the same procedure to estimate the number of genes expressed in the subsampled neurons (above threshold “3” as above). We obtained the relationship *N_genes_* = 4756 + 2127 ⋅ *log*(*N_neurons_*), and estimate a total number of to 14,920 genes expressed in the *C. elegans* nervous system.

### Transcript quantification with StringTie

For the transcript-level quantification, we used StringTie 2.2.1 using the annotation from Wormbase (WS289), without novel transcript discovery. We applied it to the aligned reads from STAR, following deduplication. Code available at: https://github.com/cengenproject/stringtie_quantif.

### Website

The output of STAR was used to generate browser tracks. First, the junction counts generated by STAR were processed, and the number of uniquely mapped reads was kept as junction count. Junctions longer than 25,000 bp, and junctions with a count lower than 3 reads were filtered out. Junction counts for individual samples were combined into neuron type average, and global sum, minimum, and maximum. The global tracks underwent a second filtering, requiring 13 and 21 reads for the maximum and sum tracks respectively. The tracks were exported to bed format using the R package rtracklayer ^88^. Second, the bam files generated by STAR were used to generate the coverage tracks using custom code, and exported to bigWig format with rtracklayer. The individual tracks can be used in JBrowse2 ^45^ or downloaded from the website. All code is available at: https://github.com/cengenproject/splicing_browser.

For the local splicing quantification, the results of the MAJIQ analysis (see above) were loaded in VOILA according to instructions (see https://majiq.biociphers.org).

For the transcript-level quantification, the quantifications from StringTie (see above) were loaded in a custom R Shiny application, source code available at: https://github.com/cengenproject/isoform_compare/.

### Binary dAS with SUPPA2

We used SUPPA 2.3 ^89^ according to the documentation. We generated all local event types (SE, SS, MX, RI, FL) with `--boundary S` based on the genome annotation (Wormbase WS289). We then quantified the event PSI by running `psiPerEvent` using the StringTie quantifications (see above) as expression file. Finally, we split the resulting TPM and PSI files by neuron type, before computing delta PSI for each pair of neuron, using the settings `-method empirical -combination -gene-correction`.

We filtered using single-cell threshold 2 ^32^ to consider an event detectable in a neuron pair only when the gene is expressed in both neurons of a pair. We considered an event differentially AS in a neuron pair if it is detectable in both neurons of the pair, has a p-value lower than 0.1 (from SUPPA with -gc option for multiple comparison), and displays a delta PSI higher than 0.3.

We used the R package GenomicFeatures ^90^ to delineate the genomic regions of interest, the package Biostrings to extract the sequence and calculate its GC content, and the conservation score from ^91^. The features were compared using a Wilcoxon test followed by Holm correction for multiple comparisons, and we applied a threshold of 0.05 to consider a comparison significant.

For microexons, we focused on cassette exons. We restricted our analysis to neuron pairs in which both neurons express the exon-containing gene (with threshold 2 above). We compared the number of exons with DAS using a two-proportion test with Yates’ continuity correction. To compare the proportion of neuron pairs with DAS, we only analyzed exons detectable in at least 10 neuron pairs (787 out of 913 cassette exons). We used a Wilcoxon test to compare the proportions. Code available at https://github.com/cengenproject/suppa_events.

### Splicing regulatory network

#### Quantification of PSI

Cassette events were extracted from the genome annotation using `suppa generateEvents` ^89^. With an approach adapted from ^92^, we then used `bedtools` and `grep` on the STAR output to count the number of inclusion reads 𝑁_𝑖_ covering the alternative exon, and the number of exclusion reads 𝑁_𝑔_ spanning the alternative splice junction. We then computed Percent Spliced-In (PSI) from normalized read counts *N̄_l_* and *N̄_e_* based on exon length *L_exon_* and read length *l_read_*:

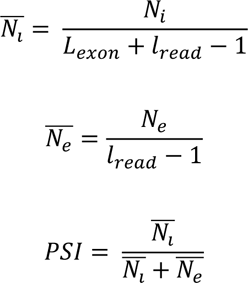

All relevant code is stored in the Github repository

https://github.com/cengenproject/quantif_exon_skipping.

Before use in the model, we removed measurements of a cassette exon in a neuron type if the exon-containing gene is not expressed in that neuron type, based on a thresholding integrating this dataset with single-cell RNA-Seq ^40^. Additionally, we only considered neuron types for which we had 3 or more biological replicates. We also filtered the cassette exons, keeping only events covered by more than 20 reads, measured in more than 70 samples from 23 neuron types, and presenting differential splicing between neuron types (standard deviation above 0.3).

#### Quantification of putative splice factor transcripts

We compiled a list of putative splice factors, available in Table S5. The transcripts are quantified using StringTie ^47^, without novel transcript discovery (see above). StringTie gives quantifications in Transcripts Per Million (TPM), which undergo a log10 transformation with a pseudocount of 1 before further processing.

#### Precision matrix estimation

Here we describe the procedure to select our network construction method (Fig S5A). We build a data matrix where the 127 rows correspond to samples, the 730 columns correspond to cassette exon PSI (172 events) or splice factor log-TPM (558 transcripts). We perform a first split: 30% of the rows (39 samples) are kept as testing set. The other 70% of samples undergo 5-fold cross-validation: each fold contains 17 or 18 samples, the training is performed on 4 folds, the validation on the held-out fold. As the splicing of a cassette exon in a neuron type can only be meaningfully measured if the gene containing that exon is expressed in that neuron, the PSI matrix contains missing values that are first imputed (using the column median or k-nearest neighbors). The training matrices *SE_train_* (containing the skipped exons) and *SF_train_* (containing the splice factors) are then transformed (using Z-score or NPN transformation), and a covariance matrix *S_trian_* is computed from the transformed values. We store the variables used for transformation (e.g. the mean and standard deviation for Z-scoring and distribution quantiles for the NPN method) for inverting the transformations later to yield predictions in the original data range. For permutation tests, the *SE_train_* values randomized within an event (i.e. within a column) after transformation, but before computing the covariance matrix. The covariance matrix is then used to estimate the precision matrix 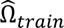 (using glasso, QUIC, CLIME, or SCIO), which is inverted to recover the estimated covariance matrix *Ŝ_train_*.

Separately, the validation fold matrices *SE_valid_* and *SF_valid_* are transformed re-using the same parameters as the training folds, to compute the covariance matrix *S_valid_*. From the estimated precision matrix, following ^55^, we extract the quadrants 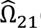(with the splice factors as rows and the cassette exons as columns), and 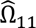 (with the cassette exons as rows and columns) and compute 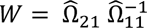, the matrix of regression coefficients. The splicing measurements in the validation set can then be estimated from the splice factors in the validation set and the precision matrix learned from the training set following:

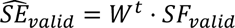

Finally, we invert the transformation of 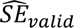 using the stored transformation variables (e.g. the mean and standard deviation for Z-scoring and distribution quantiles for the NPN method) to get back to the initial scale.

#### Model components

We tried several approaches to develop an optimal model. The data matrix *SE_train_* can be constructed from PSIs, a ratio of inclusion and exclusion counts. We reasoned that a model may have a better performance when directly estimating the inclusion and exclusion counts rather than the ratio. Thus we reconstructed counts by multiplying the PSI with the total count for that exon, and used either PSIs or reconstructed counts as the columns of *SE_train_*.

As our downstream methods are incompatible with the presence of missing values, we need to remove them from *SE_train_*. We used median imputation, where the missing value is replaced by the median of the column (i.e. the median of the cassette exon across samples). Alternatively, we used a k-nearest neighbors imputation implemented by the R package impute.

PSIs (or reconstructed counts) and log-TPMs follow very different distributions, which would distort the covariance computed by simply concatenating them. In addition, they do not follow a normal distribution, making them inappropriate for the downstream algorithms. We thus standardized the training data either with a Z-score transformation, or with nonparanormal transformations ^61^. As our cross-validation procedure requires that we transform the validation set using the parameters from the training set, and that we invert the transformation to obtain values in the original scale, we implemented these transformations in the R package projectNPN, available at https://github.com/cengenproject/projectNPN.

Finally, the estimation of the precision matrix can also be performed with several implementations. We used the R packages glasso, QUIC, FLARE (implementing CLIME) and SCIO.

#### Metrics definitions

First, we compare the covariance matrix measured in the validation set *S_valid_* to the covariance matrix estimated from the training set *Ŝ_train_*, obtained by inverting the estimated precision matrix. we focus on the quadrant containing the covariance between the skipped exons and the splice factors, as we are interested in the ability of our model to capture this relationship. We then compute the Frobenius loss as 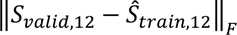 where 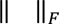 represents the Frobenius norm.

Further, we compare the skipping values measured in the validation set *SE_valid_* to the values 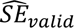 estimated from the precision matrix. To this end, we first compute the residuals: 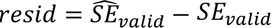. We then estimate, for each event *e*, the sum of squared residuals: 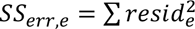 and the total sum of squares 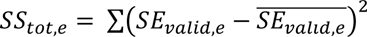 and define the fraction explained variance as 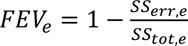. We truncate this value at 0 and average it across events to obtain the mean fraction of explained variance.

To evaluate the biological relevance of the network edges, we extract the quadrant 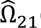 of the precision matrix (i.e. the adjacency matrix), binarize it (taking an edge for any non-zero entry), and compare it to a ground truth dataset (table S5). We compiled this dataset by a review of the literature, considering that a regulatory interaction between a splice factor and a cassette exon is “true” if a change in splicing was detected upon mutation of the splice factor. Note that this dataset suffers from several limitations, notably these interactions do not necessarily take place in the neurons we sequenced here, and these interactions may correspond to different splicing events within the same target gene. Thus, while we expect a better model to obtain a better match with this data, we do not expect a perfect match. We compute the True Positive Rate (TPR) as the fraction of interactions present in the ground truth that are captured by the model, and the False Positive Rate (FPR) as the fraction of interactions that are absent from the ground truth but reported by the model. As both TPR and FPR decrease with increased sparsity, we report the ratio TPR/FPR.

Finally, we seek to constrain the structure of the network. A very sparse network, where each splice factor has at most a single target, or a very dense network, where many splice factors have many targets, would be hard to interpret and likely not capture biologically meaningful interactions. As proposed by ^56^, we use an approximate scale-free topology criterion. For each splice factor (node in the network), we compute the degree of the node *k*, and count the number of nodes with the same degree *p(k)*. We then fit a linear regression between log(𝑘) and log(𝑝(𝑘)), and use the coefficient of determination *R*^2^ as a criterion. A high *R^2^* suggests that a power law can describe the node degrees, and that the network is scale-free.

All code related to the network modeling is available at: https://github.com/cengenproject/regression_exon_skipping

### Data and code availability

Sequencing data is available on GEO (accession GSE229078).

All code is in https://github.com/cengenproject/ (see corresponding repository names in Methods).

## Supporting information

Supplemental Table 1: samples

Supplemental Table 2: novel junctions

Supplemental Table 3: genes DAS

Supplemental Table 4: sequence comparisons

Supplemental Table 5: putative SF

Supplemental Table 6: ground truth interactions

Supplemental Table 7: metrics

Supplemental Table 8: permutations

Supplemental Table 9: adjacency matrix

## Acknowledgements

We would like to thank members of the Marc Hammarlund, Shaul Yogev, and Karla Neugebauer labs for helpful discussion and advice. The CeNGEN project is supported by NIH grant R01NS100547. E.V. is supported by grant R00MH128772. Sequencing was done with YCGA: research reported in this publication was supported by the National Institute of General Medical Sciences of the National Institutes of Health under Award Number 1S10OD030363-01A1. Some strains were provided by the CGC, which is funded by NIH Office of Research Infrastructure Programs (P40 OD010440).

## Author contributions

A.W. contributed to the acquisition, analysis, and interpretation of data and writing of the manuscript.

E.V. and A.B. contributed to the analysis of data. A.B., R.M.M., S.R.T., I.C., M.B., A.P., J.A.T., B.C. contributed to the acquisition of data. S.K., D.M.M., and M.H. contributed to the conception of the work and to writing the manuscript.

## CeNGEN Consortium members

Additional members of the CeNGEN Consortium: Cyril Cros, Berta Vidal, Maryam Majeed, Chen Wang,

Eviatar Yemini, Emily A. Bayer, HaoSheng Sun, Oliver Hobert.

## Supplementary figures

**Figure S1:**
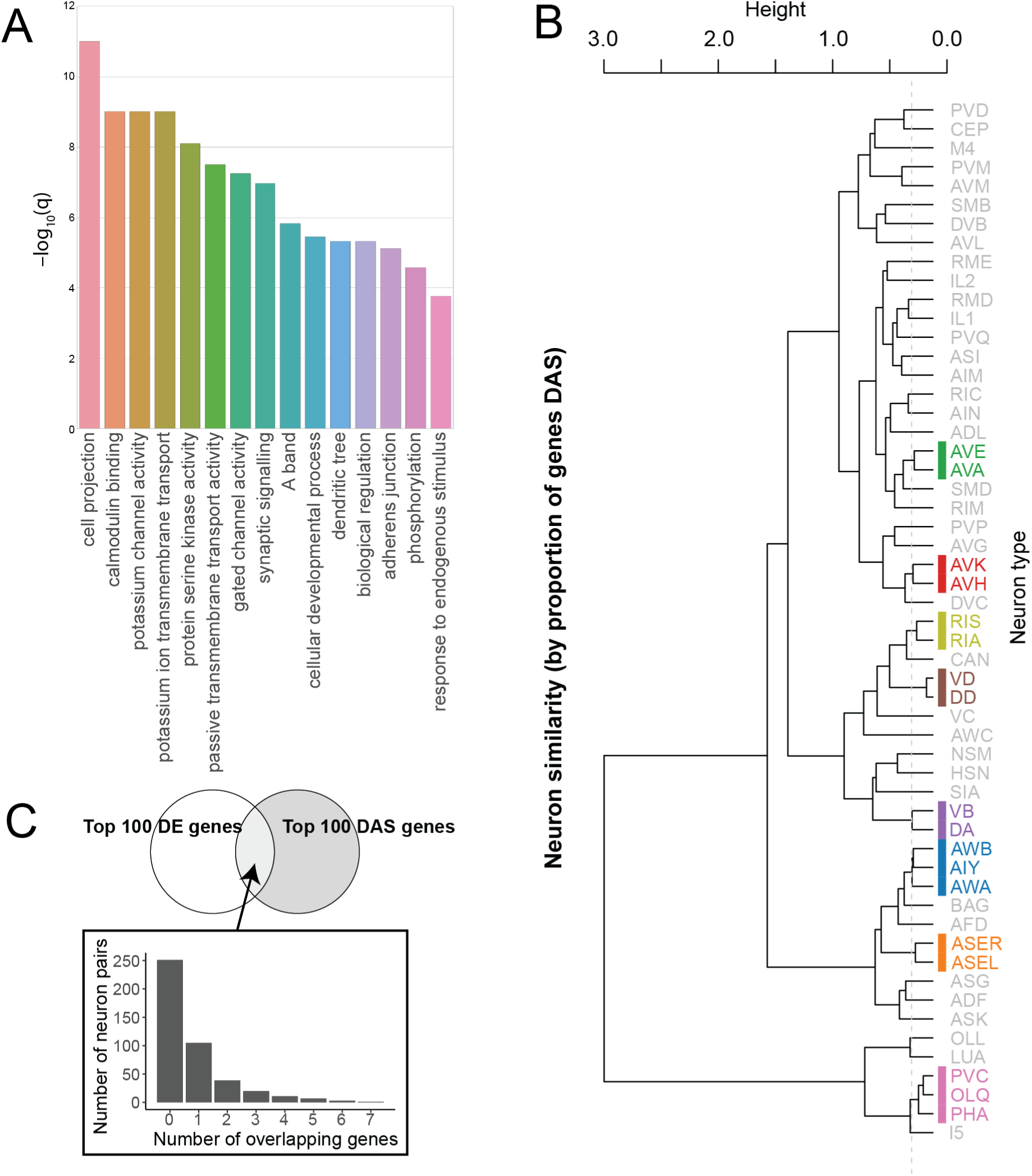
Analysis of DAS genes. **A:** Gene Ontology terms enriched among the genes with DAS in neurons. **B:** Overlap between the top 100 DAS genes and the top 100 differentially expressed genes for each neuron pair. **C:** Pairs of neurons with similar profiles, clustered by proportion of DAS genes among co-expressed genes, corresponding to the heatmap in Fig. 3C. The color blocks indicate clusters identified from a tree cut at a height of 0.2 (horizontal dashed line).

**Figure S2:**
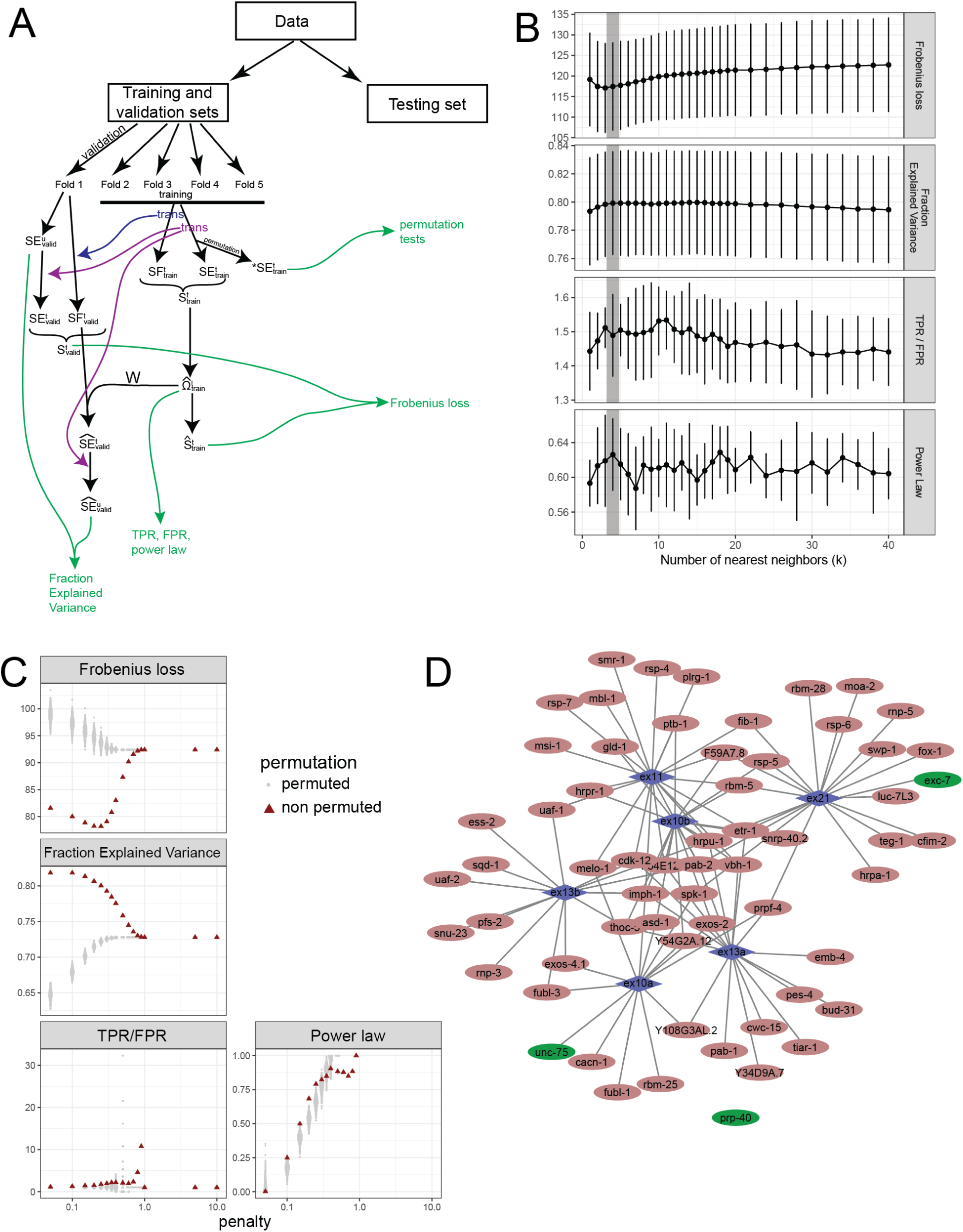
Network computation. **A:** Schematic description of the full procedure. SE: skipped exon splicing measurements, SF: splice factor expressions, S: covariance matrix, Ω: precision matrix. The superscript t or u indicates whether the matrix is transformed or untransformed, the subscript train or valid indicates whether it contains data from the training or validation folds. The green arrows and text correspond to the metrics being computed. **B:** Selection of the optimal number of nearest neighbors. The four metrics are represented at a fixed penalty of 0.2 and plotted against the number of nearest neighbors. The grey line highlights the chosen value of k=4. **C:** Metrics plotted against penalty, the red triangles indicate the computed value, the grey dots were computed after permutation of the input. **D:** Subnetwork centered on the alternative exons of unc-16. Symbols as defined in Fig. 5D.

## List of supplementary tables

S1: List of sequenced samples

S2: List of novel splice junctions

S3: List of known AS genes from literature

S4: Features of binary events that were tested, means and p-values

S5: List of putative splice factors and RBPs used for network

S6: List of “ground truth” splicing regulatory interactions

S7: Metrics for various hyperparameters during CV

S8: Metrics for permutation test on best model

S9: SF node degrees in network

